# A vaccine targeting the L9 epitope of the malaria circumsporozoite protein confers protection from blood-stage infection in a mouse challenge model

**DOI:** 10.1101/2021.11.10.465032

**Authors:** Lucie Jelínková, Yevel Flores-Garcia, Sarah Shapiro, Bryce T. Roberts, Nikolai Petrovsky, Fidel Zavala, Bryce Chackerian

## Abstract

Pre-erythrocytic malaria vaccines that induce high-titer, durable antibody responses can potentially provide protection from infection. Here, we engineered a virus-like particle (VLP)-based vaccine targeting a recently described vulnerable epitope at the N-terminus of the central repeat region of the *Plasmodium falciparum* circumsporozoite protein (CSP) that is recognized by the potently inhibitory monoclonal antibody L9 and show that immunization with L9 VLPs induces strong antibody responses that provide protection from blood-stage malaria in a mouse infection model.

## Main text

Pre-erythrocytic malaria vaccines (such as RTS,S ^1^) that largely target the immunodominant central repeat (CR) region of the *Plasmodium falciparum* circumsporozoite protein (*Pf*CSP) – a protein that densely covers the surface of invading sporozoites – provide moderate protection from human infection ^2, 3, 4^. However, the recent identification of human monoclonal antibodies (mAbs) from human volunteers immunized with an experimental irradiated whole sporozoite vaccine developed by Sanaria (*Pf*SPZ) that target epitopes in CSP outside of the CR and potently protect animal models from malaria infection have pointed to new sites of vulnerability in CSP that may be exploited using epitope-targeted vaccines ^5, 6, 7, 8, 9, 10^. We previously showed that a bacteriophage Qß-based virus-like particle (VLP) vaccine that multivalently displays a peptide representing the epitope of the CIS43 mAb, which is located at the junction between the N-terminal region of CSP and the CR ^5^, could elicit extremely durable and high-titer anti-CSP antibody responses and reduce parasite liver burden in a mouse malaria challenge model, but did not prevent blood-stage parasitemia ^10^. Here, we assessed the immunogenicity and protective efficacy of VLPs displaying the epitope of the L9 mAb, a newly described antibody that is one of the most potent anti-CSP mAbs at inhibiting parasite invasion in mouse models ^6^. The L9 epitope overlaps with the CIS43 epitope, but is centered on the minor repeat sequences at the N-terminus of the CR (Fig. 1a).

**Fig. 1:**
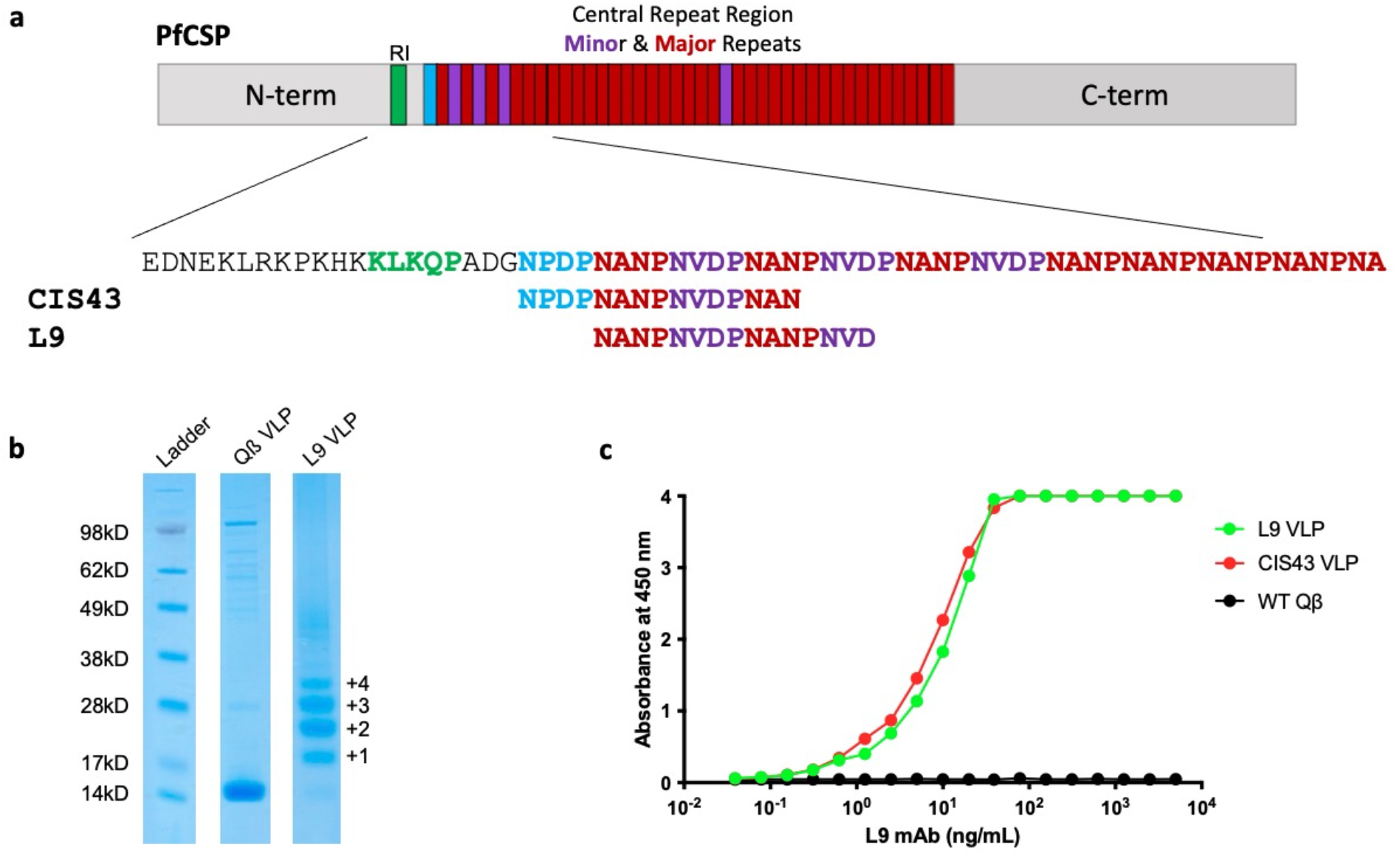
Characterization of L9 VLPs. **a** The structure of PfCSP and the location of the epitopes targeted in this study. CSP contains an N-terminal region, which contains the RI cleavage site (in green), a Junction region (between RI and the central repeat), and the central repeat (CR) Region, which contains 4 NVDP minor repeats (purple) and >35 NANP major repeats (red). The two peptide epitopes displayed on CIS43 and L9 VLPs are shown. **b** SDS-PAGE analysis of unconjugated (center lane) or L9 peptide conjugated (right lane) Qβ VLPs. The ladder of bands in the L9 VLP lane reflect individual copies of coat protein modified with 1, 2, 3, or more copies of the L9 peptide. Gel images are derived from the same experiment and were processed in parallel. Size markers are shown in the left lane. Data from a single conjugation reaction that is representative of >6 independent reactions is depicted. **c** Binding of the L9 mAb to L9 VLPs (green), CIS43 VLPs (red), or wild-type (unmodified) Qβ VLPs (black) as measured by ELISA. This experiment was performed twice, data from one representative experiment is shown.

We generated Qß VLPs that multivalently display the 15 amino acid L9 epitope (L9 VLPs) by chemically conjugating a synthetic L9 peptide to the surface of VLP using a bifunctional crosslinker. L9 VLPs display an average of 385 copies of the peptide epitope per VLP (Fig. 1b), a high valency that is associated with strong immunogenicity ^11^, and the L9 VLPs are strongly bound by the L9 mAb (Fig. 1c). L9 mAb also binds strongly to CIS43 VLPs, likely because of the overlap between the two peptide epitopes. L9 VLPs are highly immunogenic in mice; three doses of unadjuvanted L9 VLPs elicited high titer and durable anti-CSP antibodies (Fig. 2a), as was previously observed in mice immunized with CIS43 VLPs ^10^. Similar to CIS43 VLPs, L9 VLPs elicit antibodies that inhibit binding of both the L9 mAb and the CIS43 mAb to CSP (Fig. 3), indicating that induced antibodies bind to the junctional region of CSP.

**Fig. 2:**
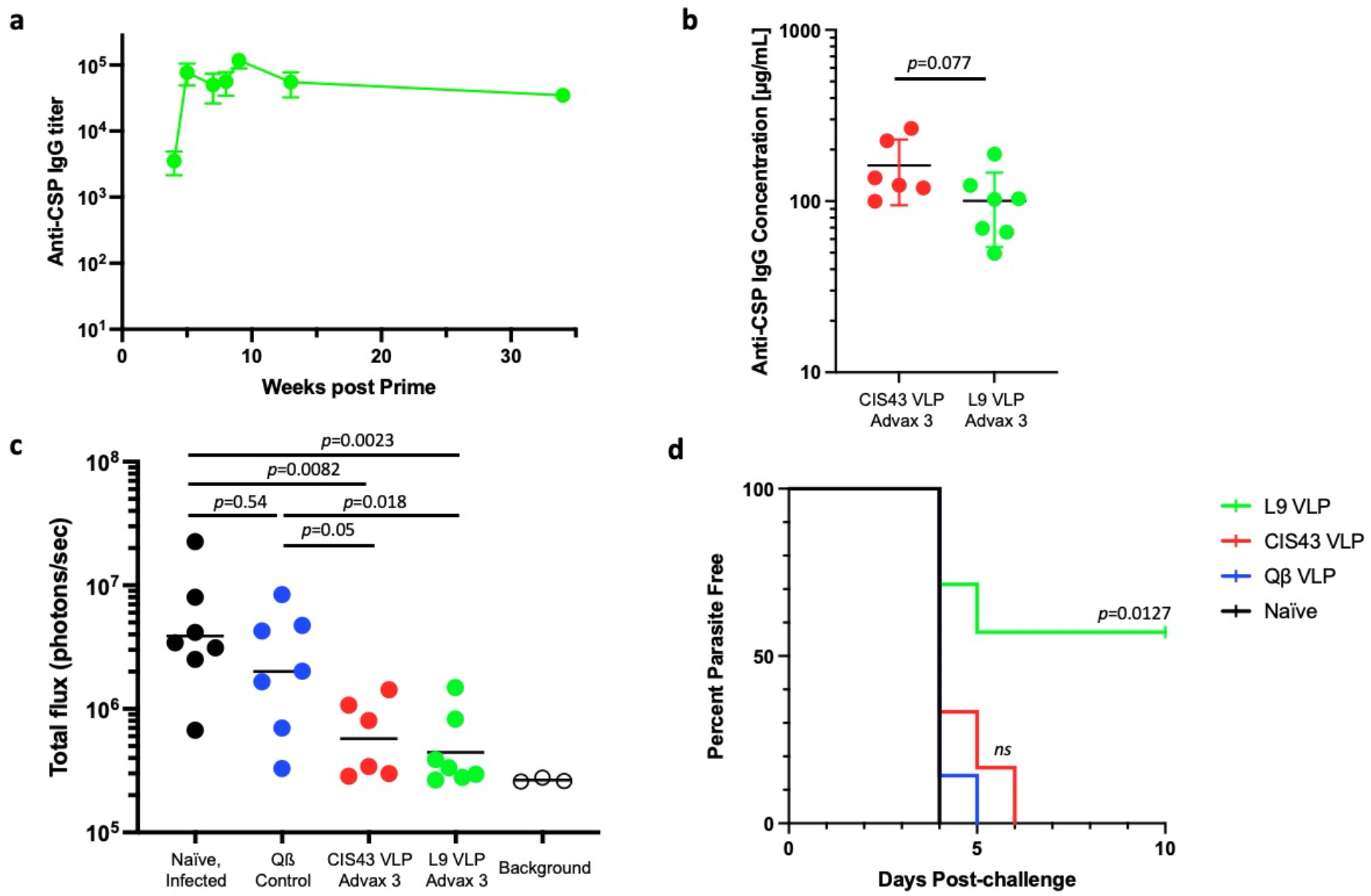
L9 VLPs elicit strong and long-lasting anti-CSP antibody responses that protect against parasitemia. **a** Mean anti-CSP IgG concentrations sampled over 34 weeks in Balb/c mice (n=6/group) immunized three times (at weeks 0, 4, & 7) with L9 VLPs without adjuvant. Error bars represent SEM. **b** Anti-CSP antibody concentrations in C57BL6 mice immunized three times with CIS43 VLPs or L9 VLPs (both with Advax-3 adjuvant; n = 6-7/group) collected 26 days following the third immunization. Antibody concentrations were compared by two-tailed t test. Serum from control (Qß VLP) immunized mice had anti-CSP antibody concentrations below the limit of detection of the assay (<0.1ng/mL). **c** Parasite liver load (as measured by luminescence) in CIS43 VLP and L9 VLP-vaccinated (or control) C57BL6 mice (n=6-7/group) following mosquito challenge. A two-tailed Mann–Whitney test was used to statistically compare groups. Background luminescence was determined using three uninfected mice. **d** Percent of parasite-free mice post-challenge as evaluated by Giemsa blood smear. Log-rank test was used to statistically compare the L9 VLP and CIS43 VLP groups to the wild-type Qß VLP control group; ns, not significant.

**Fig 3.**
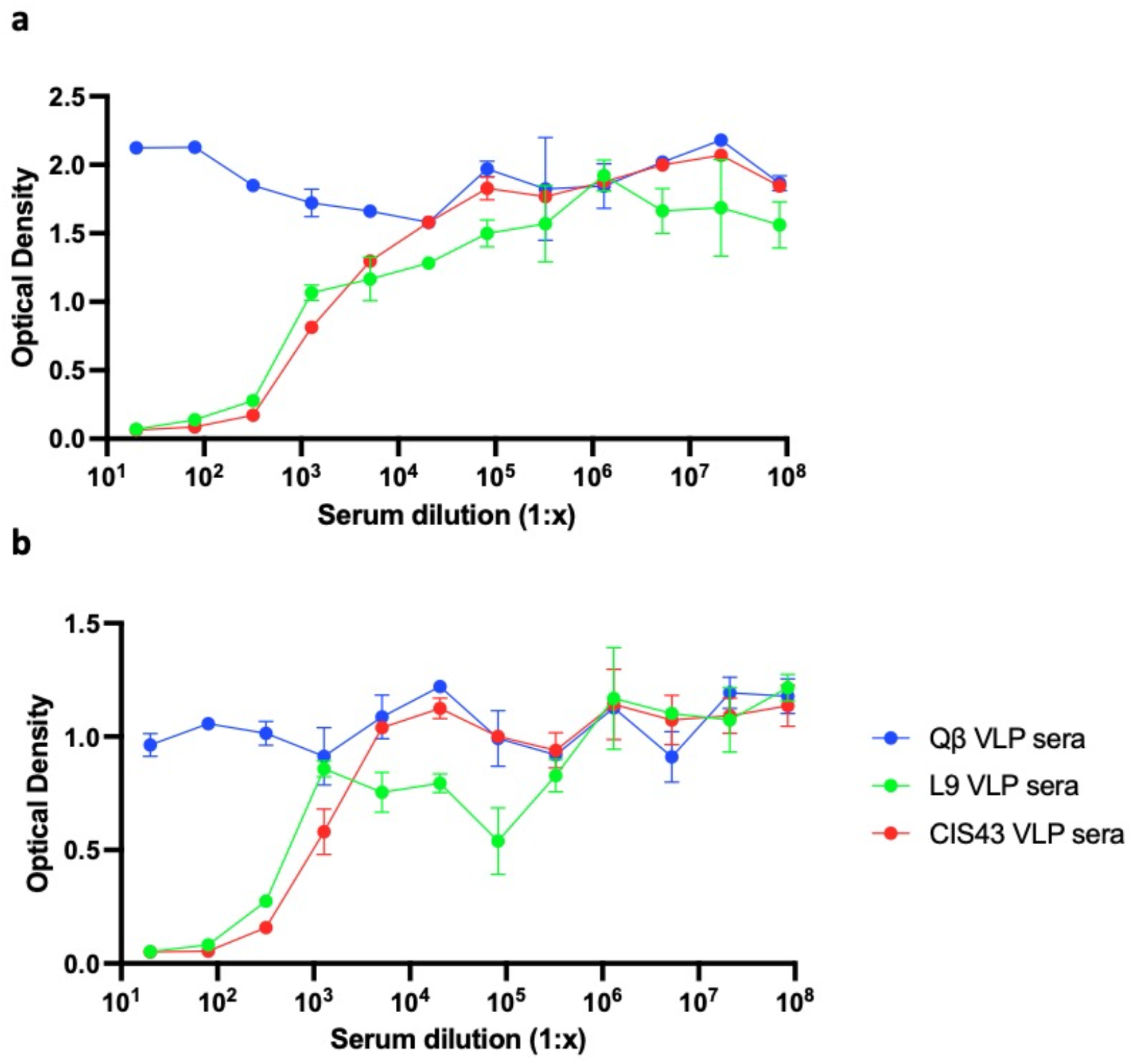
Sera from mice immunized with L9 VLPs and CIS43 VLPs inhibit the binding of mAbs L9 and CIS43 to CSP. Pooled sera from L9 VLP-immunized mice, CIS43 VLP-immunized mice, or Qß VLP-immunized mice were tested by competition ELISA for inhibition of **a** L9 mAb binding to CSP, or **b** CIS43 mAb binding to CSP. This experiment was performed in duplicate, error bars represent SEM.

To test whether L9 VLPs could confer protection from malaria challenge, C57Bl/6 mice were vaccinated with L9 VLPs, CIS43 VLPs, or, as a negative control, wild-type Qß VLPs and then challenged with malaria-infected mosquitoes. We have previously shown that co-administration of CIS43 VLPs with the adjuvant Advax-3, which is a mixture of CpG55.2 oligonucleotide (a TLR9 agonist) with aluminum hydroxide, could increase anti-CSP antibody levels, so all vaccines were adjuvanted with Advax-3. As is shown in Fig. 2b, both L9 VLPs and CIS43 VLPs mixed with Advax-3 elicited strong anti-CSP antibody responses. After three immunizations, mice were exposed to mosquitoes infected with luciferase-reporter containing transgenic *P. berghei* (*Pb*) engineered to express full-length *Pf*CSP in place of *Pb*CSP (*Pb-Pf*CSP-*Luc*) ^12^. 42 hours after challenge, liver parasite loads were measured using an intravital imaging system. Mice immunized with CIS43 VLPs and L9 VLPs had significantly lower liver parasite loads than control VLP-vaccinated mice or unvaccinated (naïve) controls (Fig. 2c). Relative to naïve mice, immunization with CIS43 VLPs reduced mean parasite loads by ~89% (similar to what we showed previously ^10^) and immunization with L9 VLPs reduced mean parasite loads by ~92%. Beginning four days after infection, blood smears from mice were evaluated for parasitemia. While all control mice and CIS43 VLP-immunized mice developed blood-stage parasitemia, 4 of the 7 mice immunized with L9 VLPs remained parasite-free 10 days after infection, indicating that these mice were protected from blood-stage infection (Fig. 2d).

Here, we evaluated and compared the efficacy of vaccines targeting two overlapping epitopes within the junctional/minor repeat regions of *Plasmodium falciparum* CSP that are recognized by the potent inhibitory monoclonal antibodies, CIS43 and L9. VLPs displaying both epitopes could elicit strong anti-CSP antibody responses and could significantly reduce parasite liver loads in experimentally challenged mice. However, only L9 VLPs could prevent blood parasitemia. The 15 amino acid CIS43 and L9 epitopes overlap by 11 amino acids, suggesting that subtle changes in the epitope targeted by anti-CSP antibodies can dramatically affect protective efficacy. It has been hypothesized that the most potent anti-CSP mAbs recognize epitopes derived from the joining of minor and major tetrapeptide repeats, including DPNA (minor/major) and NPNV (major/minor) ^6, 13, 14^. The DPNA motif is found twice in the CIS43 epitope and once in the L9 epitope, whereas NPNV is found twice in the L9 epitope and once in the CIS43 epitope. In addition, the minimal L9 binding peptide (NA<u>NPNV</u>DP) is centered on the NPNV sequence ^15^. To examine the specificity of antibodies induced by the L9 and CIS43 VLP vaccines, the binding of sera to peptides representing the L9 and CIS43 epitopes, as well as peptides representing the major repeat (NANP) and non-naturally occurring peptides representing repeat junctions (DPNA and NPNV) was assessed (Fig. 4). Both vaccines induced antibodies that bound to all five peptides, but sera from mice immunized with L9 VLPs had stronger relative binding to the L9 epitope than the CIS43 epitope, and reacted more strongly with the (NPNV)_3_ peptide than the (DPNA)_3_ peptide. In contrast, sera from CIS43-immunized mice bound more strongly to the CIS43 epitope and the (DPNA)_3_ peptide. Thus, it is possible that antibodies which more strongly target NPNV are more functionally active and, therefore, may aid in preventing blood parasitemia. Future experiments may reveal the specific nature of vulnerability of this region of CSP. Taken together, these studies indicate that L9 VLPs are a promising malaria vaccine candidate.

**Fig. 4.**
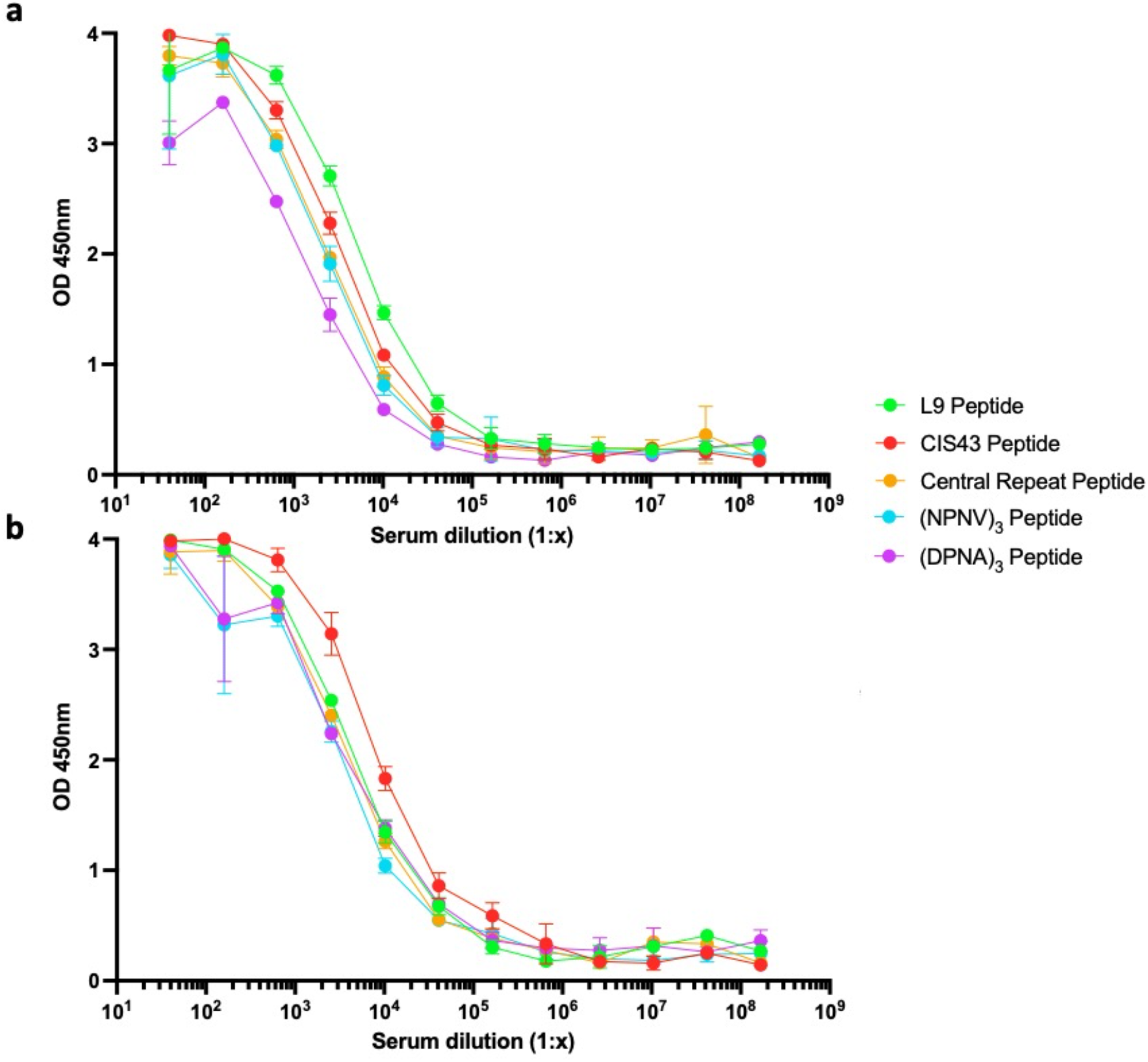
Binding of sera to specific CSP-derived epitopes. Pooled sera from **a** L9 VLP-immunized mice or, **b** CIS43 VLP-immunized mice were tested for binding to five different peptides by ELISA. This experiment was performed in triplicate, error bars represent SD.

## Methods

### Ethics

All animal research complied with the Institutional Animal Care and Use Committee of the University of New Mexico School of Medicine (Approved protocol #: 19-200870-HSC), Johns Hopkins University (Approved protocol permit #: MO18H419).

### Production and characterization of VLP-based vaccines

CIS43 VLPs were produced as described in ^10^. L9 VLPs were produced similarly; the fifteen amino acid L9 epitope peptide was synthesized (GenScript) and modified to contain a C-terminal *gly-gly-gly-cys* linker sequence (NANPNVDPNANPNVD*GGGC*) and was conjugated directly to surface lysines on Qβ bacteriophage VLPs using the bidirectional crosslinker succinimidyl 6-[(beta-maleimidopropionamido) hexanoate] (SMPH; Thermo Fisher Scientific) as previously described ^10^. The efficiency of conjugation was assessed by gel electrophoresis using a 10% SDS denaturing polyacrylamide gel followed by analysis using ImageJ software to calculate the average peptide density per VLP. Conjugation of VLPs with the L9 peptide was performed multiple times, with similar conjugation efficiencies. Presence of the L9 peptide on L9 VLPs was confirmed by ELISA. Briefly, 250ng of VLPs were used to coat wells of an ELISA plate. Wells were probed with dilutions of mAb L9 (generously provided by Robert Seder, NIH Vaccine Research Center), followed by a 1:4000 dilution of horseradish peroxidase (HRP) labeled goat anti-human IgG (Jackson Immunoresearch). The reaction was developed using 3,3’,5,5’-tetramethylbenzidine (TMB) substrate (Thermo Fisher Scientific) and stopped using 1% HCl. Reactivity was determined by measuring optical density at 450 nm (OD_450_) using an AccuSkan plate reader (Fisher Scientific).

### Mouse Immunization Studies

For the initial evaluation of immunogenicity, groups (n=6) of 4-5-week old female Balb/c mice (obtained from the Jackson Laboratory) were immunized intramuscularly with 5μg of L9 VLPs without exogenous adjuvant. Mice were boosted twice at 4 and 7 weeks after the initial prime. Challenge studies were performed using 7-8-week old C57Bl/6 mice (n=6-7/group), which are more susceptible to malaria challenge than Balb/c mice. Mice were immunized with 5μg doses of L9 VLPs, CIS43 VLPs, or wild-type control Qß VLPs in combination with 20μL of Advax-3 adjuvant (5mg/mL). Mice were immunized at days 0, 28, and 56, serum was collected at day 82, and the mice were challenged at day 84. An additional group of naïve (unimmunized) mice were included in this experiment.

### Quantitating antibody responses

Serum antibodies against full-length CSP were detected by ELISA using recombinant CSP expressed in *Pseudomonas fluorescens ^16^* (and generously provided by Gabriel Gutierrez at Leidos, Inc.) as the coating antigen, as described previously ^10^. Anti-CSP antibody concentrations were determined by generating a standard curve using known concentrations of the anti-CSP mouse mAb 2A10. Peptide ELISAs were performed as described ^10^ using peptides representing the L9 epitope (see above), the CIS43 epitope (NPDPNANPNVDPNAN), the CR major repeat (NANPNANPNANPNANPNA), and multimeric NPNV [(NPNV)_3_; NPNVNPNVNPNV] and DPNA [(DPNA)_3_; DPNADPNADPNA) peptides. All peptides were synthesized to contain a C-terminal-GGGC linker sequence. Competition ELISAs were performed by using the CSP ELISA protocol with the following modifications: after serum from L9 VLP, CIS43 VLP, or wild-type control Qß VLP immunized mice was added to the plate, 40 ng of the human mAb L9 or CIS43 (at a final concentration of 400 ng/mL) was added to each well and incubated for 30 minutes. L9 or CIS43 mAb binding to CSP was detected using HRP-labeled goat anti-human IgG at a 1:4000 dilution.

### Mouse Pb-PfCSP-Luc sporozoite mosquito challenge

Mice were challenged directly by using infected mosquitos four weeks following their third final vaccination. *Anopheles stephensi* mosquitos were infected by blood-feeding on *Pb-Pf*CSP-*Luc* infected mice. Prior to challenge, mice were anesthetized with 2% Avertin, and then exposed to six mosquitos for a blood meal for 10 minutes. Following feeding, the number of mosquitos positive for a blood meal was determined. Liver luminescence was assessed 42 hours post-challenge by intraperitoneally injecting anesthetized mice with 100 μl D-luciferin (30mg/ml) and then determining liver luminescence using an IVIS Spectrum Imaging System (Perkin Elmer). Beginning four days after challenge, blood smears were collected daily and then evaluated by Giemsa staining for parasitemia.

## Data Availability

The datasets used and/or analyzed in the current study are available from the corresponding author upon reasonable request.

## Acknowledgements

This study was supported, in part, by a generous contribution to the UNM Foundation in honor of Jeffrey Michael Gorvetzian in support of biomedical research excellence at the University of New Mexico School of Medicine and by T32-AI007538 (to L.J.). F.Z. thanks the Bloomberg Philanthropies for continued support. Development of Advax adjuvants was supported by funding from National Institute of Allergy and Infectious Diseases of the National Institutes of Health under Contracts HHS-N272201400053C, HHS-N272200800039C and U01-AI061142.

## Author Contributions

L.J., F.Z., N.P. and B.C. designed the study and drafted the manuscript. L.J. engineered the VLPs, performed the mouse experiments at the University of New Mexico, performed the analysis of immune responses, and analyzed all data. Y.G-F., S.S. and F.Z. performed the mouse challenge experiments and assisted in the statistical analysis of the data resulting from those experiments. B.T.R. performed peptide ELISA experiments and assisted with the revision of the manuscript. N.P. supplied adjuvants.

